# Antinociceptive and Antioxidant Activities of Methanolic Extract of Leaves of *Azadirachta Indica* (Neem)

**DOI:** 10.1101/694505

**Authors:** Akash S Mali, Mahesh B Thorat, Dhairyasheel M. Ghadge, Kumodini A. Nikam, Shraddha Sawant, Farida Shaikh, Nayana Vhatkar, Swapnja Shinde, Kavyashree Gonghade, Munaf A. Tamboli

## Abstract

**Graphical abstract:** 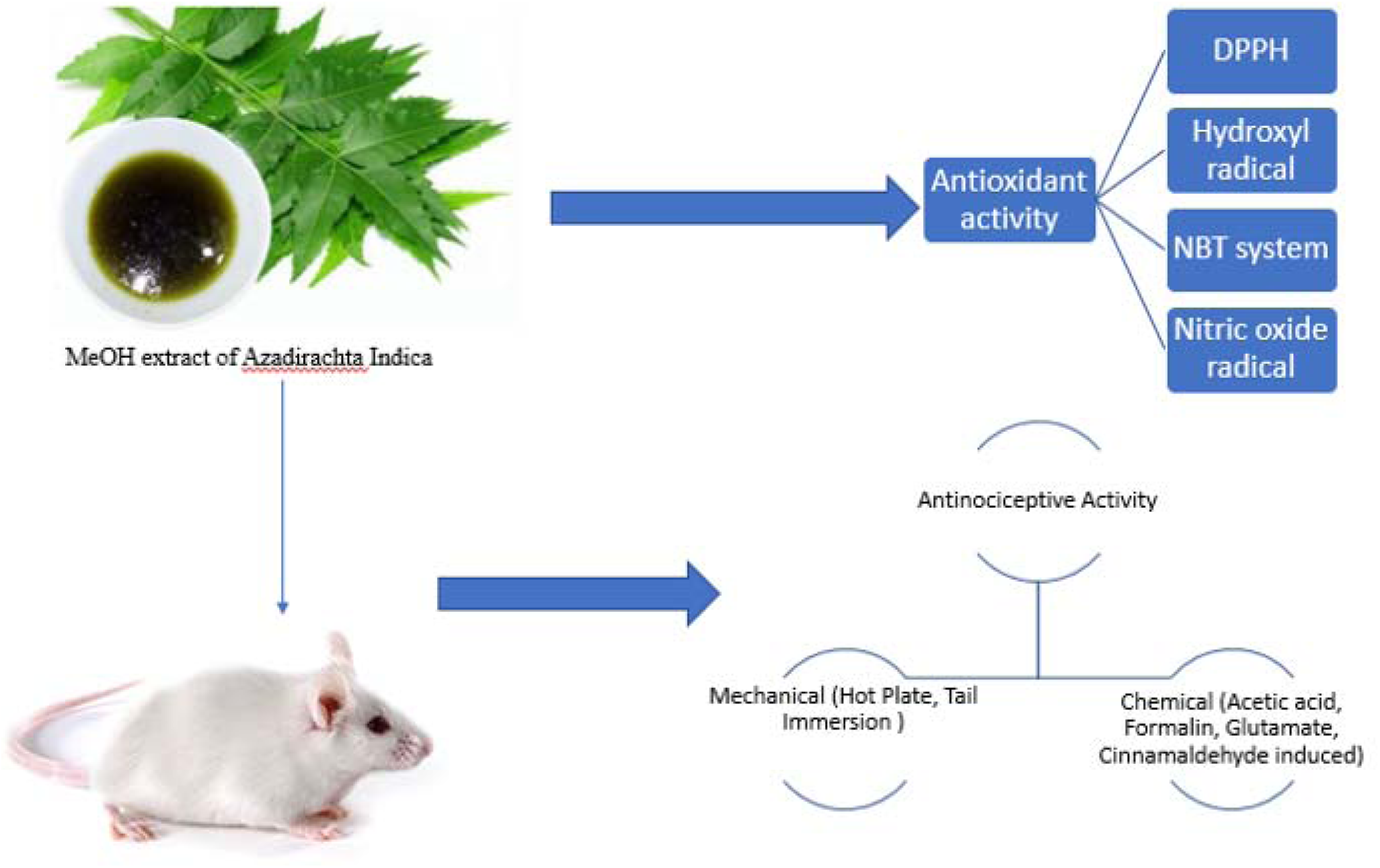

**Background:** *Azadirachta indica* (Neem) is a communal plant of *Meliaceae* family called Neem Or *Kadunimb* in Maharashtra, India Neem stated anti-inflammatory through regulation of proinflammatory enzyme activities with *COX* and *LOX* enzyme. Previous studies show that Azadirachta Indica (neem) and its chief constituents play essential role in anticancer management via the modulation of different molecular pathways including NF-κB, p53, PI3K/Akt, Bcl-2, pTEN and VEGF. Many parts of the plant are traditionally used in the treatment of various pharmacological action, the analgesic activity of Neem Seed Oil has already reported but Neem Leaves.

**Methods:** The antinociceptive activity of Azadirachta Indica Leaves (AZIL) was examined using heat-induced-mechanical (hot-plate and tail-immersion test) and chemical-induced (acetic acid, formalin, glutamic acid, cinnamaldehyde) nociception models in mice at 50,100, and 200 mg/kg doses. ATP-sensitive K+ channel pathway, cyclic guanosine monophosphate (cGMP) pathway and involvement of opioid system was also tested using glibenclamide, methylene blue and naloxone/morphine respectively. The methanolic extract of leaves of *A.Indica* was assessed by using different *in vitro* antioxidant models of screening like scavenging of 1,1-diphenyl-2-picryl hydrazyl (DPPH) radical, nitric oxide radical, superoxide anion radical, and hydroxyl radical.

**Results:** AZIL showed antinociceptive activity and antioxidant activity. In both hot plate and tail immersion tests AZIL significantly increases the latency to the thermal stimuli. In acetic acid-induced writhing test the extract repressed the number of abdominal writhing. Similarly, AZIL produced substantial dose-dependent inhibition of paw licking in both neurogenic and inflammatory pain induced by intraplanar injection of formalin. As well, AZIL also expressively withdrawn cinnamaldehyde-induced pain and the glutamate-induced pain in mice. It was also proved that pretreatment with naloxone significantly reversed the antinociception produced by AZIL in mechanical tests signifying the involvement of opioid system in its effect. Furthermore, administration of methylene blue, enhanced AZIL induced antinociception while glibenclamide, an ATP-sensitive K+ channel antagonist, could not converse antinociceptive activity induced by AZIL.

**Conclusion:** Based on the results of the present study it can be said that AZIL keeps significant antinociceptive activity which acts in both central and peripheral mechanisms.

**Ethnopharmacological relevance:** Azadiracta Indica having family Melliases is one of the common medicinal plant in United states and south Asia including India, Pakistan, Bangladesh. Various parts of the plants used in treatment of many inflammatory conditions, skin diseases, tooth protection, antidiabetic in the form of oil and herbal porridge.

## Introduction

*Azadirachta Indica* (Neem) is an evergreen, fast growing ancient tree, generally known as Miracle tree. Neem has an excellent of medicinal properties and active ingredients like *azadirachtin, nimbidin, flavonoids, alkaloids, saponins, tannins* which play vital role in various pharmacological actions [1]. *Azadirachta indica* has become significant in the global context today because it offers answers to the key concerns facing mankind. Since ancient time man is in search of remedies for pain. Pain is the commonest symptom that takes a patient to the hospital. Traditional medicine provides a substitute through which this quest can be fulfilled. Neem is a fast-growing perennial tree, is a native of Indian subcontinent, Africa, America. It is acknowledged for over 4000 years now and is called ‘arishtha’ in Sanskrit meaning ‘perfect, reliever of sickness, hence it has been acclaimed as ‘Sarbarogaribarini’ [2]. This miracle tree’ Neem also exhibits antibacterial, antiviral, antioxidant, antimutagenic, immunomodulatory, anti-inflammatory, antihyperglycemic, antiulcer, antimalarial, antifungal, and anticarcinogenic properties [2] Chemical constitutes which is demanded to possess analgesic and anti-inflammatory properties. NSAID’s and opioids are used nowadays, is limited by its own side effects. The analgesic activity using the Neem Seed Oil (NSO) has already been done but not the *Azadirachta indica* Leaf Extract (AZIL) [2,3]. Hence the present study is done to evaluate the analgesic effect of AZIL on albino mice as well determine antioxidant properties of AZIL via different models.

### Chemicals

All chemicals and solvents were of analytical grade and obtained from Loba Chemicals Ltd, Mumbai. 1,1-diphenyl,2-picryl hydrazyl (DPPH) was obtained from Sigma, USA. The other chemicals used were sodium nitroprusside, sulphanilamide, *O*-phosphoric acid, nachyl ethylene diamine dihydrochloride, glacial acetic acid, nitroblue tetrazolium (NBT), ethylene diamine tetra acetic acid (EDTA), riboflavin, and Fe-EDTA. morphine sulphate, diclofenac sodium, naloxone (Lobachem,Mumbai), trichloroacetic acid (TCA), methanol, formalin, cinnamaldehyde, methylene blue, L-glutamic acid (Merck, Germany), glibenclemide, and DMSO (Lobachem, Mumbai). UV Visible spectrophotometer (Shimadzu 1700) was used for recording the spectra.

### Experimental animals

Swiss albino mice (20–25 g) of either sex were obtained from National Institute of Bioscience (1091/GO/bt/S/07/CPCSEA), Dist-Pune (Maharashtra) and were adapted for 10 days under standard housing conditions (23°±2°C; 50-55% RH with 12:12 h light/dark cycle). approved by Committee for the Purpose of Control and Supervision on Experiments on Animals (CPCSEA). The animals had free access to mice food (Lipton Gold Mohr, India) and water. The animals were habituated to laboratory conditions for one week prior to the experimental protocol to minimize any nonspecific stress. The experimental protocol was approved by the Institutional Animal Ethics Committee by Gourishankar Institute of Pharmaceutical education and research Limb Satara, India (Approval no. GIPER/2017-18/09).

### Plant material and extract preparation

The leaves of *A. Indica* were collected from Satara region, Maharashtra, India in November 2017. The collected samples were then identified by Botany Department, Yashwantrao Chavan College of Science, Satara, India. Powdered dried leaves (50.75g) were soaked with 250 ml of methanol with occasional stirring at 23±2°C for 72 Hrs. The extract was then filtered with the help of sterilized cotton filter and Buchner funnel. The solvent was removed by rotary evaporator and 20 g extract was obtained (Yield 39.40%). This crude extract was used for the further studies.

### Phytochemical screening

AZIL was qualitatively tested for the detection of carbohydrates, saponins, flavonoids, tannins, alkaloids, glycosides, reducing sugars by standard procedures (See Supplement).

## *In vitro* antioxidant activity

### DPPH assay

To 1 ml extract of different concentrations, 1 ml solution of DPPH (0.1 mM) was added. An equal amount of MeOH and DPPH solution served as control. After 20 mins of incubation in the dark, absorbance was measured at 517 nm.1g Ascorbic acid was used as standard. The experiment was done in triplicate and the percentage scavenging was calculated. [4]

### Scavenging of nitric oxide radical

Nitric oxide was produced from sodium nitroprusside and measured by Griess reaction. [9,10] Sodium nitroprusside (5mM) in standard phosphate buffer saline solution (0.75 mM, pH 7.4) was incubated with different concentrations of (25-250 μg/ml) of the methanolic extract dissolved in phosphate buffer saline after that the tubes were incubated at 25°C. after 5 Hrs 0.5 ml of solution was removed and diluted with 0.5 ml of Griess reagent (2g of 1% sulphanilamide, 2 g of 0.1% napthyl ethylene diamine dihydrochloride and 5 ml of 2% *O*-phosphoric acid). The absorbance was read at 546 nm. Ascorbic acid was used as standard. The experiment was done in triplicate. [5]

### Hydroxyl radical scavenging activity

AZIL with different concentrations (25-250μg/ml) were taken in different test tubes and evaporated on water bath.1 ml DMSO, 0.5 ml of EDTA, and 1 ml Fe-EDTA were added and the reaction was started by adding 0.5 ml ascorbic acid to each of the test tubes. Test tubes were capped firmly and heated in water bath at 75-95°C for 20 m. after that the reaction was completed by addition of 1 ml of ice-cold TCA (17.5%, w/v) in every test tube and kept at RT for 5 to 10 min. The formaldehyde was measured by adding 2.5 ml Nash’s reagent. This reaction mixture was kept aside for 15 min for color development. [4,5] Intensity of yellow color was measured spectrophotometrically at 412 nm. Ascorbic acid was used as standard.

### Scavenging of superoxide radical by riboflavin-NBT system

The assay was constructed on the capacity of the sample to inhibition blue formazon formation by scavenging the superoxide radicals generated in the riboflavin-NBT system. The reaction mixture contains 50 mM phosphate buffer pH 7.6, 20 g riboflavin, 12 mM NBT. Reaction was started by enlightening the test samples of the extract (25-250μg/ ml). The absorbance was measured at 590 nm [6,7].

### Drugs and treatments

The control group orally administered by deionized water (0.1 mL/mice) 30 min before the experiments. The positive control group intraperitoneally received standard drug morphine in hot plate, and tail immersion test at the dose of 2 mg/kg acetyl salicylic acid (100mg/kg) in acetic acid-induced writhing and diclofenac sodium (10 mg/kg) in formalin induced licking, glutamate-induced paw licking, cinnamaldehyde-induced licking test 15 min before the experiments. AZIL was administered orally at the doses of 50, 100, and 200 mg/kg 30 min before the experiments. To assess the involvement of opioid-mediated antinociceptive activity, naloxone was administered (at the dose of 1 mg/kg 10 min before morphine sulfate (2mg/kg) or AZIL (50, 100, and 200 mg/kg) running in the hot plate and tail immersion test. Methylene blue (15 mg/kg) and glibenclamide (5 mg/kg) were intraperitoneally injected 15 min before control administration to evaluate the involvement of cyclic guanosine monophosphate(cGMP) and ATP-sensitive K+ channel pathway respectively. All the doses of drugs and AZIL were prepared using ddH_2_O.

## Antinociceptive analysis

### Hot plate test

The mice that showed fore paw licking, withdrawal response within 15s on hot plate (55± 0.5°C). Mice were fasted overnight with water given ad libitum. The animals were treated with morphine or AZIL and were placed on Eddy’s hot plate kept at a temperature of 55 ± 0.5°C. A cut off period of 20s was maintained to avoid paw tissue damage [8]. The response in the form of fore paw licking, withdrawal of the paw(s) or jumping was recorded on 30, 60, 90, and 120 min following treatment.

### Tail immersion test

To evaluate the central analgesic property the tail immersion test was performed. This method is based on the observation that morphine like drugs prolongs the tail withdrawal time from hot water in mice [8,9]. 1-2 cm of tail of the mice pretreated with morphine or AZIL were immersed in warm water kept constant at 50±0.5°C. The latency between tail submersion and refraction of tail was recorded. A latency period of 20s was maintained to avoid tail tissue damage in mice. The latency period of the tail-withdrawal response was taken, as the index of antinociception and was determined at 30, 60, 90, and 120 min after the administration of morphine or AZIL.

### Acetic acid-induced writhing test

Acetic acid-induced writhing test was performed to evaluate the peripheral antinociceptive activity of AZIL in chemical-induced pain. The mice were treated with drug or AZIL and then the writhing was induced by injecting 0.5% acetic acid after 10 and 20 min, respectively, at the dose 10 ml/kg body weight. Five min after the injection of acetic acid, the mice were observed, and the number of writhing was counted for 30 min [9,10]. The contractions of the abdomen, elongation of the body, twisting of the trunk and/or pelvis ending with the extension of the limbs were considered as complete writhing.

### Glutamate-induced nociception

20 μ L of glutamate (10 μM/paw) was injected into the ventral surface of the right hind paw of mice 30 min after AZIL treatment and 15 min after injection of diclofenac sodium. The mice were observed for 10 min following glutamate injection and number of paw licking was counted as an indication of nociception [11].

### Formalin-induced nociception

Mice were injected with 10 μl of a 2.5% formalin solution (0.92% formaldehyde) made up in saline into the subplantar region of the right hind paw 60 min after AZIL treatment and 10 min after injection of diclofenac sodium. Licking of the injected paw was recorded as nociceptive response from 0-5 min (neurogenic phase) and 10-20 min (inflammatory phase) after formalin injection [10,11].

### Cinnamaldehyde-induced nociception

10 μl of cinnamaldehyde (10 nmol/paw prepared in saline)were injected intraplantarly in the ventral surface of the right hind paw. Animals were observed individually for 5 min following cinnamaldehyde injection. The amount of time spent licking the injected paw was recorded with achronometer and was considered as indicative of nociception. The animals were treated with AZIL orally 1 hr before cinnamaldehyde injection. Control animals received vehicle (10 ml/kg orally) [12,13].

## Analysis of the possible mechanism of action of AZIL

### Involvement of opioid system

The possible participation of the opioid system in the antinociceptive effect of AZIL was examined by injecting naloxone hydrochloride (2 mg/kg i.p.), a non-selective opioid receptor antagonist, 15 min prior to the administration of either morphine or AZIL. Then the hot plate and tail immersion latencies were measured at 30, 60, 90 and 120 min with the same cut off time of 20 s for the safety of animals [14].

### Involvement of cyclic guanosine monophosphate (cGMP) pathway

To verify the possible involvement of cGMP pathway in the antinociceptive action caused by AZIL the mice were pretreated with methylene blue (20 mg/kg), a non-specific inhibitor of Nitric oxide/guanylyl cyclase, intraperitonially 15 min before administration of ether diclofenac sodium or AZIL. Then the nociceptive responses against 0.6% acetic acid injection were observed for 30 min, starting from5 min post injection. The numbers of abdominal writhing were counted as indication of pain behavior [16].

### Involvement of ATP-sensitive K+ channel pathway

Possible contribution of K+ channel in the antinociceptive effect of AZIL was evaluated by previously described method [16,17]. The mice were pre-treated with glibenclamide (10mg/kg), an ATP-sensitive K+ channel inhibitor, intraperitonially 15 min before administrative of either diclofenac Sodium or AZIL. Following the injection of acetic acid, the animals were immediately placed in a Perspex chamber and the number of writhing was recorded for 30 min, starting from 5 min post injection.

### Statistical analysis

The results are presented as MEAN±SEM. The statistical analysis of the results was performed using one-way analysis of variance (ANOVA) followed by Dunnett’s test or Bonferroni’s test as appropriate using SPSS 25.0 software. Differences between groups were considered significant at a level of p<0.001 and p< 0.05. The results of the tail immersion and hot plate tests were given with percentage of the maximal possible effect (%MPE), which was calculated using the following formula.

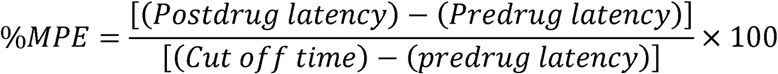

## Results

### Phytochemical screening

The preliminary screening revealed the presence of alkaloid, carbohydrate, glycoside, steroid, flavonoid, saponin and tannin in AZIL (Table no.1).

**Table No. 1.** Phytochemical analysis of AZIL.

| Sr.No | Phytochemicals | Water | Methanol |
| --- | --- | --- | --- |
| 1 | Alkaloids | ++ | +++ |
| 2 | Flavonoids | ++ | +++ |
| 3 | Saponins | + | + |
| 4 | Tannins | + | ++ |
| 5 | Reducing sugars | ++ | +++ |
| 6 | Glycosides | ++ | +++ |
| 7 | Phenol | + | ++ |
| 8 | Terpanoids | - | + |
| +:Low; ++: Medium; +++: High, -: Absent |  |  |  |

### Acute toxicity

From the acute toxicity test it has been found that the LD_50_ of AZIL is 758.58 mg/kg in mice.

### Antioxidant activity

Six concentrations ranging from 0.05 to 1 mg/ml of the methanolic extract of *A. indica* were tested for their antioxidant potential using different *in vitro* models. Study indicate that free radicals were scavenged by test compounds at different concentrations (0.05,0.1,0.2,0.4,0.6,0.8,1.0 mg/ml). The maximum inhibitory concentration (IC_50_) in nitric oxide radical, DPPH, superoxide, hydroxyl radical scavenging activity, IC_50_ were found to be 6.38, 6.65, 9.21,and 4.35 mg/ml, respectively. The antioxidant model and percentage scavenging of each concentration of extract and standard are shown in Table 2.

**Table No. 2.** Antioxidant activity of Azadiracta Indica.

| Sr.No | Antioxident models | % RSA of AA 200mg/ml (Control) | Azadiracta indica extract (mg/ml) |  |  |  |  |  |  |
| --- | --- | --- | --- | --- | --- | --- | --- | --- | --- |
|  |  |  | 0.05 | 0.1 | 0.2 | 0.4 | 0.8 | 1.0 | IC<sub>50</sub> |
| 1 | DPPH* | 76.35 | 29.35 | 34.15 | 47.65 | 49.32 | 53.647 | 61.25 | 07.65 |
| 2 | Nitric Oxide | 78.65 | 41.36 | 44.36 | 51.26 | 69.35 | 70.32 | 74.36 | 06.38 |
| 3 | Hydroxyl radicle | 61.37 | 37.65 | 47.62 | 52.36 | 56.65 | 61.26 | 73.95 | 04.35 |
| 4 | Superoxide radicle | 83.65 | 43.65 | 51.65 | 61.65 | 67.96 | 74.64 | 84.36 | 09.21 |

AA- Ascorbic acid, \*-1,1-diphenyl, 2-picryl hydrazyl

### Hot plate test

The antinociceptive effect of AZIL and morphine in hot plate test are given in Table 3. AZIL at 100 and 200 mg/kg doses significantly increased the reaction time to the thermal stimulus (p<0.05). The antinociceptive effect was dose-dependent as we observed stronger effect at 200 mg/kg dose than 50 mg/kg dose. Morphine showed highest latency at all the observation periods. The extract also showed significant increase in latency to the thermal stimuli at 50, 100, and 200 mg/kg doses (p<0.05). Naloxone exerted significant (p<0.05) antagonistic effect on the antinociceptive activity of AZIL and morphine.

**Table No. 3.** Effect of AZIL extract in hot plate test.

| Treatment | Dose (mg/kg) | Latency period (Seconds) (%MPE) |  |  |  |  |
| --- | --- | --- | --- | --- | --- | --- |
|  |  | Pre-treatment | 30 Min | 60 Min | 90 Min | 120 Min |
| Control | 0.1ml/mice | 9.1 ±0.35 | 10.25±1.1* | 12.65±0.6 | 14.68±0.7 | 14.99±2.6** |
| Morphine | 2 | 10.2 ±0.10 | 9.91±0.67<br>(-15.15) | 12.78±0.12<br>(5.18) | 15.54±0.51<br>(6.69) | 15.98±0.09<br>(7.48) |
| Nalaxone(NLX) | 1 | 11.68 ±0.8 | 11.31±1.1 | 10.87±1.3 | 9.84±0.98 | 9.03±0.21 |
| Control+NLX | 1 | 11.35 ± 0.3 | 9.97±1.54 | 9.57±0.12 | 11.65±0.68 | 12.08±1.5 |
| Morphine+NLX | 2+1 | 11.98 ± 0.2 | 12.36±0.54<br>(2.10) | 12.78±0.51<br>(1.66) | 13.14±1.3<br>(1.48) | 13.57±0.65<br>(1.47) |
| AZIL | 50 | 9.54 ± 0.22 | 9.99±0.25<br>(2.25) | 10.24±0.24<br>(1.40) | 12.35±0.78<br>(3.51) | 12.96±0.67<br>(3.12) |
|  | 100 | 10.35 ± 0.6 | 10.94±0.58<br>(3.00) | 12.25±0.59<br>(2.38) | 13.65±1.4<br>(4.14) | 14.85±0.60<br>(4.10) |
|  | 200 | 10.96 ± 0.7 | 11.58±0.24<br>(5.3) | 12.22±0.14<br>(2.56) | 13.52±0.11<br>*(3.23) | 15.17±0.65<br>(3.86) |
| AZIL+NLX | 50+1 | 11.61 ± 0.1 | 11.72±0.12<br>(0.59) | 11.97±0.25<br>(0.74) | 12.03±0.84<br>(0.53) | 12.68±0.27<br>(0.98) |
|  | 100+1 | 12.13 ±0.54 | 12.43±0.30<br>(1.67) | 12.97±0.68<br>(1.75) | 13.37±0.44<br>(1.59) | 13.96±0.29<br>(1.69) |
|  | 200+1 | 13.03 ± 0.5 | 13.12 ± 0.21<br>(0.53) | 13.26±0.32<br>(0.48) | 13.54±0.56<br>(0.66) | 13.87±0.11<br>(0.79) |

Each value is presented as the mean ± SEM (n = 5). AZIL =Azadiracta Indica Leaves extract; NLX = Naloxone.
\*p < 0.05 compared with the control group (Dunnett's test).

### Tail immersion test

The antinociceptive activity of AZIL and morphine in tail immersion test has been given in Table 4. AZIL at all three doses (50,100, and 200 mg/kg) significantly increased the latency period to hot-water induced thermal stimuli (p<0.001) in a dose-dependent manner. Morphine showed highest latency, however, the extract also showed significant latency at 50, 100, and 200 mg/kg doses (p< 0.001) at different observation time. Naloxone exerted significant (p < 0.05) antagonistic effect on the antinociceptive activity of AZIL at all three doses and of morphine throughout the observation periods. (Table 4)

**Table No. 4.** Effect of AZIL extract in tail immersion test.

| Treatment | Dose (mg/kg) | Latency period (Seconds) (%MPE) |  |  |  |  |
| --- | --- | --- | --- | --- | --- | --- |
|  |  | Pre-treatment | 30 Min | 60 Min | 90 Min | 120 Min |
| Control | 0.1ml/mice | 3.39±0.22 | 3.68±0.2 | 3.72±1.3 | 3.81±0.65 | 3.89±0.80 |
| Morphine | 2 | 4.11±0.1 | 5.84±0.9<br>(6.68) | 7.84±0.2<br>(6.67) | 8.63±0.54*<br>(4.71) | 6.95±0.5*<br>(2.45) |
| Nalaxone(NLX) | 1 | 5.51±0.5 | 4.98±0.1 | 4.54±0.64 | 4.11±0.25 | 3.95±0.67 |
| Control+NLX | 1 | 4.21±0.5 | 4.65±0.51 | 4.98±0.68 | 5.67±0.21* | 5.87±0.63 |
| Morphine+NLX | 2+1 | 3.97±0.54 | 4.14±0.41<br>(0.58) | 4.36±0.84<br>(0.66) | 4.57±0.51<br>(0.60) | 4.21±0.09*<br>(0.20) |
|  | 50 | 3.33±0.23 | 3.87±0.81 | 4.25±0.10 | 5.11±0.2* | 5.00±0.61 |
| AZIL |  |  | (2.02) | (1.62) | (5.89) | (1.43) |
|  | 100 | 3.61±0.5 | 4.02±0.24<br>(1.55) | 5.21±0.58<br>(2.83) | 6.21±0.87<br>*(3.00) | 5.31±0.82<br>(1.46) |
|  | 200 | 3.84±0.43* | 4.51±0.29<br>(2.56) | 5.98±0.16<br>(3.81) | 6.44±0.17<br>(4.01) | 5.64±0.26<br>(1.54) |
| AZIL+NLX | 50+1 | 3.29±0.55 | 3.61±0.18<br>(1.19) | 3.89±0.27<br>(1.05) | 3.94±0.67<br>(0.7) | 4.06±0.95<br>(0.65) |
|  | 100+1 | 3.86±0.14 | 3.72±0.34<br>(-0.53) | 3.44±0.68<br>(-0.72) | 3.94±0.24*<br>(0.09) | 3.11±0.19<br>(-0.64) |
|  | 200+1 | 2.62±0.87 | 2.55±0.21<br>(-0.25) | 2.43±0.41<br>(-0.33) | 2.96±0.54*<br>(0.41) | 2.21±0.12<br>(-0.34) |

Each value is presented as the mean ± SEM (n = 5). AZIL =Azadiracta Indica Leaves extract; NLX = Naloxone.
\*p < 0.05 compared with the control group (Dunnett's test).

### Acetic acid-induced writhing test

The effect of oral administration of AZIL using the abdominal constriction test in mice is shown in table 5. It was found that AZIL was able to inhibit the nociceptive effects induced by acetic acid compared to the control group (Deionized water) at the doses of 50, 100, and 200 mg/kg, respectively (p<0.001). The percentage inhibition of constrictions was calculated as 69.94% (Acetyl salicylic acid,100 mg/kg), 18.55% (AZIL, 50mg/kg), 41.95% (AZIL, 100 mg/kg), and 50% (AZIL, 200 mg/kg).

**Table No. 5.** Effect of AZIL on acetic acid-induced writhing in Mice.

| Sr no | Treatment | Dose(mg/kg) | No of Writhings | Inhibition(%) |
| --- | --- | --- | --- | --- |
| 1 | Control | 0.1 ml/mice | 37.3 ± 3.2 | - |
| 2 | Acetyl salicylic acid | 100 | 11.21± 2.1 | 69.94 |
| 3 | AZIL | 50 | 30.58± 1.3* | 18.55 |
|  |  | 100 | 21.65± 1.1* | 41.95 |
|  |  | 200 | 18.65± 1.8** | 50.00 |

\*P < 0.05, \*\*P < 0.01 vs. control, [Values are mean ± SE from six animals in each group.]
\*p < 0.001 compared with the control group (Dunnett's test).

**Table No. 6.** Effect of AZIL in glutamate-induced nociception.

| Sr no | Treatment | Dose(mg/kg) | Licking Time (Sec) | Inhibition(%) |
| --- | --- | --- | --- | --- |
| 1 | Control | 0.1 ml/mice | 121.6 ± 4.2 | - |
| 2 | Diclofenac Sodium | 10 | 39.65± 3.1 | 67.39 |
| 3 | AZIL | 50 | 107.00± 1.3** | 12.00 |
|  |  | 100 | 64.36± 1.1* | 47.07 |
|  |  | 200 | 51.23± 1.8* | 57.87 |

\*P < 0.05, \*\*P < 0.01 vs. control, [Values are mean ± SE from six animals in each group.]
\*p < 0.001 compared with the control group (Dunnett's test).

### Formalin test

AZIL produced a dose-related inhibition of formalin induced nociception and caused inhibition of both neurogenic (0–10 min) and inflammatory (10–30 min) phases of formalin-induced licking test at the doses of 50, 100, and 200 mg/kg when compared with control group (Deionized water) (Table 7). Antinociceptive effect was more pronounced in the second phase of this model of pain. Diclofenac sodium (10 mg/kg, i.p.) reduced formalin induced nociception in both phases (p < 0.001).

**Table No. 7.**
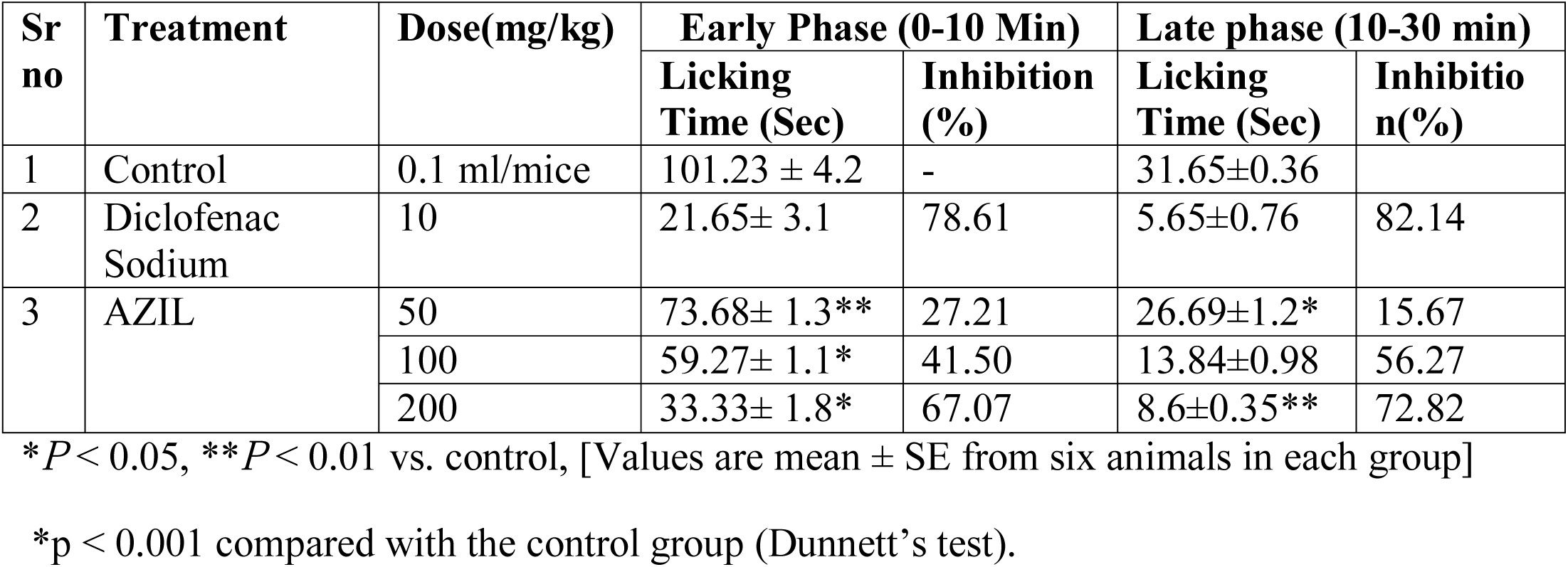
Effect of AZIL in formalin-induced nociception.

### Glutamate-induced nociception

The antinociceptive activity induced by oral administration of AZIL was dose-dependent. It showed that AZIL at the doses of 50, 100, and 200 mg/kg produced significant prohibition of the glutamate-induced nociception test (Table 6). Diclofenac sodium (10 mg/kg) was used as a standard drug, which showed 82.14 % inhibition of licking as compared to the control group. All treatments displayed significant antinociceptive activity compared together with the control group (Deionized water).

### Cinnamaldehyde-induced nociception

The result of cinnamaldehyde-induced nociception showed that administration of AZIL at 50, 100, and 200mg/kg dose produced dose-dependent inhibition of the cinnamaldehyde-induced neurogenic nociception with the percentage of inhibition of 30.37%, 65.18% and 72.68%, respectively (Table 8). Only 50 mg/ kg AZIL treatment showed no significant difference compared with the control group, at the same time as the rest presented noteworthy antinociceptive activity (p < 0.001).

**Table No. 8.** Effect of AZIL on cinnamaldehyde-induced nociception.

| Sr no | Treatment | Dose(mg/kg) | Licking Time (Sec) | Inhibition(%) |
| --- | --- | --- | --- | --- |
| 1 | Control | 0.1 ml/mice | 62.05 ± 0.65 | - |
| 2 | Diclofenac Sodium | 10 | 9.6± 0.10* | 83.92 |
| 3 | AZIL | 50 | 43.2± 1.3** | 30.37 |
|  |  | 100 | 21.6± 0.35 | 65.18 |
|  |  | 200 | 16.95± 1.8* | 72.68 |

\*P < 0.05, \*\*P < 0.01 vs. control, [Values are mean ± SE from six animals in each group.]
\*p < 0.001 compared with the control group (Dunnett's test).

### Involvement of cyclic guanosine monophosphate (cGMP) Pathway

The study showed at the effects of 50, 100, and 200 mg/ kg AZIL and methylene blue (20 mg/kg) treatments. Methylene blue administration expressively inhibited acetic acid-induced abdominal writhing (Fig. 1). Given together, methylene blue significantly increased AZIL (200 mg/kg) induced antinociception compared to the control group.

**Figure No 1 and 2.**
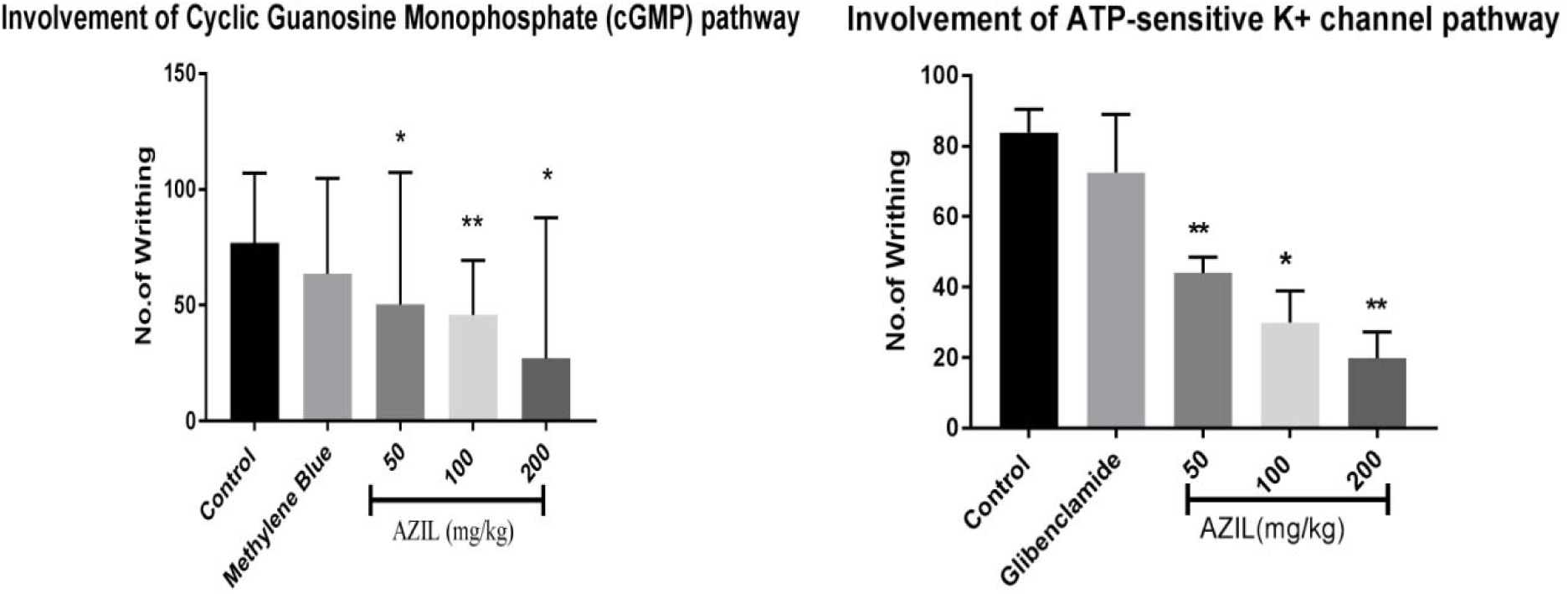
Mechanism of Study. Effects of AZIL on involvement of cyclic guanosine monophosphate (cGMP) pathway and ATP-sensitive K+ channel pathway. Values are presented as mean ± SEM (n = 5). **p < 0.001 compared with the control group (ANOVA followed by post hoc Dunnett’s test)

### Involvement of ATP-sensitive K+ channels pathway

This study appeared at the effects of 50, 100, and 200 mg/ kg AZIL and glibenclamide (10 mg/kg) treatments. It was found out that glibenclamide administration alone did not alter abdominal writhing count when assessed through the injection of 0.6% acetic acid (Fig. 2). When given together, the antinociceptive activity of AZIL was noticeably decreased by glibenclamide at the doses of 50,100 and 200 mg/kg, respectively.

This study elaborates that oral administration of AZIL produce a potent and dose-dependent antinociceptive effect in mechanical induced and chemical nociception models even AZIL acts centrally and peripherally. The important increase in latency time in hot plate by AZIL (p< 0.05) at different doses submits the central antinociceptive activity of AZIL. The effect experiential in the tail immersion test, which is particularly for centrally acting analgesics, carried out by hot plate test [17]. To find out the possible mechanism of action of AZIL we analyzed the effect of naloxone, a non-selective opioid receptor antagonist Vs the antinociceptive effect of AZIL. The reverse effect was determinative against morphine in both the hot plate and tail immersion test.

These studies confirm that the antinociceptive effect of AZIL may occur through opioid receptors at the spinal and supraspinal level. Some studies reveled mu and delta opioid receptors are involved in spinal mechanism, and mu 1 and 2-opioid receptors may mediate mainly supraspinal analgesia [18]. However, it can be predicted that the central antinociceptive effect of AZIL may be protuberant on mu-opioid receptors. Peripheral antinociceptive activity evaluated by acetic acid induced writhing test. Due to releasing pain mediators writhing test is globally accepted as a model visceral pain. Oral administration of AZIL produced significant decrease in acetic acid-induced writhing. IP administration of acetic acid increases the level of bradykinin cyclooxygenase, Interleukin 1,8 beta, lipoxygenase, prostaglandins, histamines, TNF-alpha in the peripheral tissue fluid [18]. The inhibition of writhing response maintains the peripheral antinociceptive effect of AZIL [15]. In formalin test AZIL presented important antinociceptive activity in both neurogenic (early phase) and inflammatory (late phase). Formalin-induced pain is constantly inhibited by analgesic and anti-inflammatory drugs like morphine, diclofenac sodium. This study shows AZIL significantly decreases the cinnamaldehyde-induced pain, which probably involved with TRPA1 receptor located in C-fibers reducing the pain [15,18]. Presence of alkaloids, glycosides, steroids, carbohydrates, saponins, tannins and flavonoids revealed by Preliminary phytochemical screening. Flavonoids suppress the intracellular calcium level elevation, as well as the release of pro-inflammatory mediators (TNF alfa, IL 1 beta). In future our laboratory interested to find out role of flavonoids in antinociceptive mechanism. [22]. In addition, we investigate the involvement of cGMP pathway in the antinociceptive effect of AZIL. Nociceptive activity is depending on activation or deactivation of cGMP [19,20]. Intracellular cGMP concentrations are regulated by the action of guanylyl cyclase and by the rate of degradation by cGMP-specific phosphodiesterase. Nitric oxide elevates level of cGMP by activation of soluble GCs, which impacts physiological functions including pain and analgesia [21]. cGMP acts on the ion channels directly or through the activation of protein kinases and phosphodiesterase [23,24]. Future study includes determination of involved pathways in nociception which might helpful to provide therapeutic strategy in disease condition and provide appropriate use of medicinal plant.

## Conclusions

It can be concluded that AZIL possesses important antinociceptive activity in both chemical and heat induced pain models (mechanical) in mice. The antinociceptive effect of AZIL is most likely mediated via inhibition of peripheral mediators and central inhibitory mechanisms. These results support the traditional use of this plant in different painful conditions. Further investigations are required to perceive the mechanisms of action of AZIL and to identify the active constituents that may be used as a lead compound for new drug development.

## Acknowledgements

The authors are grateful to Professor Dr. Yadav A.V., Principal, Gourishanakar Institute of Pharmaceutical education Research limb Satara, for his permission to use the facilities of the Pharmacology and Pharmaceutical chemistry Laboratory.

## Ethics approval

All the experimental mice were treated following the Ethical Principles and Guidelines from The Committee for the Purpose of Control and Supervision of Experiments on Animals (CPCSEA) The Institutional Animal Ethical Committee (1988/PO/Re/S/17/CPCSEA) of Gourishankar Institute of Pharmaceutical education and Research approved all experimental rules.

